# Network Rerouting Under Ayahuasca: Temporally and Hemisphere-Resolved EEG Connectomics

**DOI:** 10.64898/2025.12.08.693032

**Authors:** Caroline L. Alves, Fernanda Palhano-Fontes, Thaise G. L. de O. Toutain, Loriz Francisco Sallum, Christiane Thielemann, Draulio Barros de Araujo

## Abstract

Ayahuasca profoundly alters conscious experience, yet robust, time-resolved EEG markers of its network-level effects remain limited. We combined machine learning with complex-network analysis to quantify how functional connectivity reorganizes across time and hemispheres in resting-state EEG from a randomized, double-blind, placebo-controlled trial including three 5-min sessions: pre-dose (T1), 2 h post-dose (T2), and 4 h post-dose (T3). The cohort consisted of naïve ayahuasca users, a population known to exhibit attenuated or more stable acute responses, making the detection of network-level changes particularly challenging. Connectivity was estimated using multiple metrics and sliding windows (10–120 s), and network features were computed and averaged to ensure statistical validity. A representation-selection step identified Spearman correlation and an intermediate temporal scale as optimal, with classification performance peaking at 60–70 s (independent-test AUC and accuracy = 0.93). Linear mixed models revealed a bilateral decrease in eigenvector centrality (weaker hub influence), increased degree heterogeneity in the right hemisphere, and reduced global efficiency in the left. Edge-level analyses localized these effects: Posterior-left connections weakened acutely (lowest at T2), whereas right temporal–central coupling transiently strengthened (highest at T2). Together, these convergent results support a mechanistic summary: as hub-centric short-cuts weaken, communication is increasingly routed through alternative, more distributed—and less efficient—pathways, with a right-lateralized expression at a later time. Methodologically, the window-optimized, hemisphere-resolved, and edge-validated pipeline extends prior EEG work and highlights temporal scale (approximately 60 s) as a biologically meaningful parameter for detecting psychedelic-induced network reorganization.

## 1 Introduction

Used for centuries by Indigenous peoples of the Amazon basin in ceremonial and healing contexts, ayahuasca is made by combining N,N-dimethyltryptamine (DMT)–containing leaves with MAO-A–inhibiting *β*-carboline alkaloids that enable DMT’s oral activity, which induces significant effects on consciousness and perception [1].

Recent neuroimaging studies have begun to elucidate how ayahuasca influences brain network dynamics, particularly focusing on alterations in functional connectivity and activity within key neural networks. For example, a key study [2] demonstrated that ayahuasca intake leads to a significant decrease in activity in the core regions of the Default Mode Network (DMN), including the posterior cingulate cortex (PCC) and medial prefrontal cortex (mPFC). Furthermore, functional connectivity within the PCC/precuneus was notably reduced after ayahuasca ingestion. These findings indicate a relationship between the modulation of DMN activity and connectivity and the altered state of consciousness associated with ayahuasca.

According to the entropic brain hypothesis in [3], ayahuasca ingestion has been associated with an increase in Shannon entropy in brain functional networks, indicating increased unpredictability and complexity in neural activity. This increase in entropy aligns with subjective reports of “mind expansion” during psychedelic experiences.

A systematic review [1] of human neuroimaging studies on ayahuasca highlighted consistent modulation of large-scale neural networks, particularly within the DMN and SN. These findings underscore that ayahuasca reliably affects brain regions involved in self-referential and interoceptive processing.

Building on this evidence, individual studies have further characterized these network changes. For example, Palhano-Fontes et al. [2] reported acute reductions in PCC/precuneus connectivity within the DMN shortly after ayahuasca intake. In contrast, Pasquini et al. [4] described subacute effects, including increased connectivity in the anterior cingulate cortex within the SN and decreased connectivity in the PCC within the DMN. Notably, this study also detected enhanced SN–DMN coupling one day after the ayahuasca session, with these alterations associated with changes in interoceptive and affective processing.

Although ayahuasca contains the psychoactive compound DMT, the majority of mechanistic neuroimaging studies have focused on isolated DMT rather than the full ayahuasca brew. As a result, the temporal and network-level characteristics of ayahuasca—which unfold more slowly and last longer than the rapid, short-lived effects of inhaled or intravenous DMT [5]—remain comparatively less explored. Unlike the rapid and short-lived effects of inhaled or intravenous DMT, ayahuasca produces a slower and more prolonged alteration of brain activity [5]. Findings from DMT research offer important insights about the neural mechanisms that may also contribute to ayahuasca’s effects. For example, complementing fMRI reports of DMN/SN modulation, our earlier work showed that EEG investigation within the DMT cohort consistently implicated two electrode pairs—TP8–C3 and FC5–P8; regarding the network metrics, the results indicated concurrent decreases in integration and segregation with DMT [6]. Integration denotes the brain’s capacity for efficient, global information exchange across distributed regions (typically associated with shorter network paths and stronger inter-module communication), whereas segregation denotes the organization of processing into specialized modules with dense intra-module coupling and relatively sparse bridges between modules. More studies using DMT further contextualize these effects. Timmermann et al. [7] found that global functional connectivity increased while resting-state network integrity and segregation decreased. In a separate study, Timmermann et al. [8] found robust increases in spontaneous signal diversity accompanied by pronounced alpha/beta power decreases—consistent with a more entropic and less constrained neural regime. Alamia et al. [9] showed that suppressed top-down (alpha-band) backward waves together with enhanced forward waves reflect weakened top-down integrative signaling. Complementing these findings, Singleton et al. [10] demonstrated reductions in global control energy during DMT, indicating lower constraints for transitions between brain states—compatible with reduced integrative demands.

In our previous ayahuasca EEG cohort described in Alves et al. [11], we observed that the effects were concentrated in frontal and temporal regions, with F3–PO4 emerging as a new marker. In that dataset, community-level organization expanded and network-wide communication slowed, indicating decreased integration.

### 1.1 Gap and aims

Despite convergent fMRI and EEG evidence that ayahuasca and DMT alter large-scale brain organization (Table 1), several important gaps remain. First, the temporal scale at which ayahuasca most prominently reorganizes functional connectivity has not been clearly established. Second, the hemispheric and circuit-level specificity of these effects remains insufficiently mapped. Third, the balance between integration and segregation across the acute post-dose window has not been characterized in a time-resolved manner. Finally, although previous studies have documented network alterations following ayahuasca, most investigations have been performed in naïve participants, making it unclear whether consistent temporal and spatial patterns emerge across different analytic approaches.

**Table 1.** Summary of key findings from ayahuasca and DMT studies relevant to large-scale brain network organization and EEG markers. **Symbols:** ↑ increase; ↓ decrease. Upper rows (1-5) = ayahuasca-focused; lower rows (6-9) = DMT-focused.

| Study | Sample Size | Key Findings |
| --- | --- | --- |
| [2] | 9 experienced users | ↓ DMN (PCC/mPFC); ↓ PCC–precuneus FC. |
| [4] | 43 naïve users (22 ayahuasca; 21 placebo) | ↑ SN (ACC); ↓ DMN (PCC); ↑ SN–DMN after 24 h. |
| [3] | 7 experienced users | ↑ Shannon entropy; ↓ global integration. |
| [1] | Systematic review | Consistent DMN/SN modulation. |
| [11] | 16 participants | Frontal/temporal focus; F3–PO4; ↓ integration. |
| [6] | 35 participants | TP8–C3, FC5–P8; ↓ integration and segregation. |
| [8] | 13 participants | ↓ $\alpha$ and $\beta$ power; ↑ signal diversity. |
| [7] | 20 participants | ↑ global FC; ↓ integrity/segregation. |
| [10] | 14 participants | ↓ global control energy. |

**Abbreviations:**
- DMN—Default Mode Network. - SN—Salience Network. - PCC—posterior cingulate cortex. - mPFC—medial prefrontal cortex. - ACC—anterior cingulate cortex. - FC—functional connectivity. - EEG—electroencephalography. - fMRI—functional magnetic resonance imaging. - DMT—N,N-dimethyltryptamine.

Accordingly, this study pursues three main aims, evaluated specifically in a cohort of ayahuasca-naïve participants:

#### Aim 1. Time-resolved network reorganization

Determine how ayahuasca modulates the balance between large-scale integration (efficient global exchange) and segregation (specialized modular processing) across the pre-dose, ∼2 h, and ∼4 h states, using a machine-learning–guided, optimized connectivity representation.

#### Aim 2. Hemispheric and regional specificity

Test for lateralization and identify the cortical territories most implicated, as suggested by prior work—particularly frontal/temporal regions and posterior-left vs. right temporal-central couplings—leveraging the optimized connectivity representation.

#### Aim 3. Edge-level mapping

identify the specific inter-regional connections whose connectivity changes robustly under ayahuasca, summarizing them into a concise edge-level map that explains the observed network reorganization.

## 2 Methods

To provide an overview of the analytical workflow, Figure 1 summarizes the complete methodological pipeline adopted in this study—from EEG preprocessing to feature extraction and regional network analysis. Each stage is detailed in the following subsections.

**Fig. 1.**
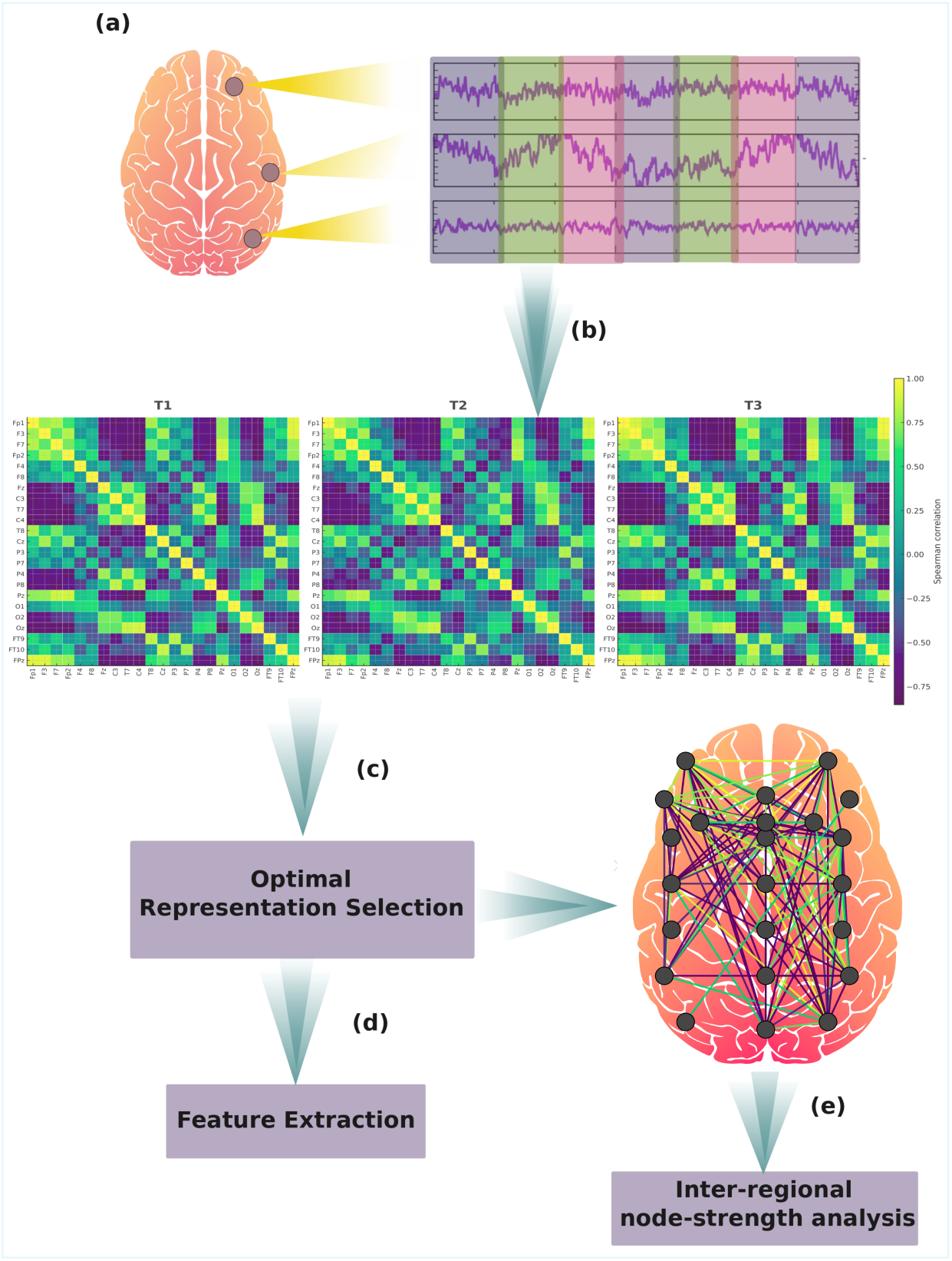
Summary of the methodological framework used in this study. (a) *EEG preprocessing:* Raw EEG data were filtered, artifact-corrected, and segmented as described in subsection 2.1. Cleaned time series were then divided into multiple sliding windows; the colored blocks (purple, green, and pink) illustrate consecutive 5-minute segments used as input for connectivity estimation. (b) *Connectivity matrix generation:* For each window, functional connectivity matrices were computed using multiple correlation– and information-theoretic metrics (subsection 2.2). The matrices shown in the figure correspond to the Spearman representation using a 60-second window, identified as optimal by the machine-learning pipeline. Matrices were computed for three time points: pre-dose (T1), ∼2 h post-dose (T2), and ∼4 h post-dose (T3). (c) *Machine-learning framework:* A supervised ML pipeline was used to identify the combination of correlation metric and temporal resolution that provided the most informative representation of ayahuasca-induced brain dynamics (subsection 2.3). (d) *Feature extraction and averaging:* From the selected optimal representations, 26 complex-network measures describing integration, segregation, efficiency, modularity, and topological complexity were computed across windows and averaged to obtain one representative feature vector per subject, condition, and time point (subsection 2.4). (e) *Interregional node-strength analysis:* Using the optimal representation, region-to-region connectivity strengths were computed to characterize hemispheric and temporal changes at the anatomical-network level (subsection 2.5). T1 = pre-dose; T2 ≈ 2 h post-dose; T3 ≈ 4 h post-dose.

### 2.1 Dataset and preprocessing

We analyzed EEG data obtained from a randomized, double-blind, placebo-controlled, parallel-arm clinical trial conducted by the Clinical Psychedelic Research Unit at Onofre Lopes University Hospital (HUOL), as described in [12]. The study included 50 healthy adults (aged 18–60) with no prior experience with ayahuasca. Ethical approval was granted by the HUOL Research Ethics Committee (protocol # 579,479) and the trial was registered on ClinicalTrials.gov (NCT02914769). EEG data were collected by members of the original clinical trial team, which includes co-authors of the present study; for this analysis, we used the recordings shared with us with appropriate permission.

Participants received a single oral dose of either ayahuasca or placebo (1 mL/kg). The ayahuasca contained, on average, 0.36 mg/mL of N,N-dimethyltryptamine (DMT), corresponding to a dose of approximately 0.36 mg/kg, which is considered a moderate to high dose based on previous clinical studies. The placebo was designed to replicate the sensory properties of ayahuasca without psychoactive effects.

Resting-state EEG was recorded in a controlled, low-stimulation environment at three time points: before dosing (0h) and approximately two and four hours after administration. Each session lasted five minutes, during which participants rested quietly with their eyes closed. For this study, we extracted five-minute EEG segments from each of these three time points. Data from participants in the ayahuasca group were labeled as T1 (pre-dose), T2 (2h post-dose), and T3 (4h post-dose). Corresponding time points for the placebo group were labeled as P1, P2, and P3, respectively.

We applied an independent preprocessing pipeline using MATLAB and EEGLAB. Signals were band-pass filtered between 1 and 50 Hz. Non-cortical channels (M1, M2, E1, E2, EMG1–3, ECG1–2) were excluded, and the remaining 23 scalp electrodes were re-referenced to the average. EEG data were segmented into 1-second epochs. Epochs containing artifacts exceeding ±150 µV were automatically rejected, followed by manual inspection.

For the ayahuasca group, two participants were excluded due to poor data quality, resulting in a final sample of 23 subjects (T1–T3) included in the connectivity and network analyses.

For the placebo group, several recordings at P1 and P2 showed signs of drowsiness or loss of wakefulness, which compromised signal stability and prevented a reliable mapping of temporal changes across the three sessions. However, all placebo participants provided usable EEG data at P3. Because P3 corresponds to the only fully awake, high-quality placebo condition, these recordings were retained exclusively for the machine-learning analyses, where they served as a comparison condition to support model training and validation. All subsequent graph-theoretical and node-strength analyses focused solely on the ayahuasca group (T1–T3), since only this group provided consistent, artifact-free data over the full time series.

After preprocessing, the cleaned five-minute EEG segments from each retained time point (T1–T3 for the ayahuasca group and P3 for the placebo group) were used to generate multiple connectivity matrices, varying both correlation metric and temporal resolution. A machine learning framework was then applied to identify which of these matrices provided the most informative representation of brain connectivity (see subsection 2.3). Only after this selection step, the optimal matrices were transformed into graphs and used for complex network feature extraction, as described in subsection 2.4. Furthermore, we also performed an inter-regional node-strength analysis based on the full five-minute segments, as detailed in subsection 2.5.

### 2.2 Connectivity Matrix Generation

In this study, we implemented an extended and novel pipeline specifically designed to capture the temporal dynamics of EEG connectivity in the context of ayahuasca administration. We adopted a methodology consistent with our previous work [6, 11, 13–20], combining complex network theory and machine learning to characterize functional brain connectivity. This extension represents a key methodological innovation of the present work.

To explore hemispheric and time-resolved patterns, we computed functional connectivity matrices using five distinct methods: Spearman’s correlation (SC) [21], Pearson’s correlation (PC) [22], Canonical Correlation Analysis (CCA) [23], Ledoit-Wolf shrinkage covariance (LW) [24], and Transfer Entropy (TE) [25]. These matrices were constructed independently for three electrode configurations: the full set of scalp electrodes and the left and right hemispheres.

A central novelty of this study compared to our previous work lies in the generation of connectivity matrices across multiple temporal resolutions. For the ayahuasca group, connectivity matrices were derived from each of the three valid time points (T1, T2, T3), using sliding windows of 10, 20, 30, 40, 50, 60, 70, 80, 90, 100, 110, and 120 seconds, as well as the entire 5-minute segment (hereafter referred to as the *full segment*).

For the placebo group, only the P3 recordings were retained due to wakefulness instability at P1 and P2. Accordingly, matrices for the placebo group were generated only for P3, and these were used exclusively within the machine-learning pipeline to provide a control condition for model training and evaluation.

Importantly, only matrices derived from sliding windows were labeled with a time suffix (e.g. _10, _20, *…*, _120), whereas matrices computed from the full segment remained unsuffixed.

### 2.3 Machine Learning Framework for Optimal Representation Selection

In this study, machine learning (ML) was not employed as the primary analytical tool but rather as a systematic strategy to identify the most informative representation of brain connectivity. Specifically, our objective was to determine (i) which correlation metric best captured functional brain changes induced by ayahuasca and (ii) whether different temporal resolutions of the connectivity matrices provided additional discriminative power.

We employed a hold-out validation scheme, splitting the data into 75% training and 25% testing partitions with stratification by group and time point. To prevent data leakage, all sliding-window representations (10–120 s) were generated after the train/test split. Within the training set, a stratified 10-fold cross-validation combined with a grid search method, commonly used in the literature [26–30], was used to identify the optimal combination of correlation metric and temporal resolution while tuning model hyperparameters. Model evaluation at both stages was based on multiple performance metrics, including the area under the curve (AUC) [29, 31–33], accuracy [31, 34–37], recall, and precision [38–41].

Once the best-performing correlation metric and window size were identified and evaluated with Support Vector Machines (SVM) [42], we employed a second machine learning step to evaluate whether the selected representations could effectively differentiate between brain states across time points. For this validation, we trained and evaluated multiple algorithms—multilayer perceptron (MLP) [43], random forest (RF) [44], and logistic regression (LR) [45].

Importantly, the purpose of this machine learning framework was not to develop a predictive model per se, but to guide the choice of connectivity representation that most effectively characterizes the brain dynamics under ayahuasca. This ensured that subsequent complex network analyses were grounded in the most informative and reliable representations of the EEG data. The results of this selection process—including the comparative performance of each connectivity metric and temporal resolution—are presented in Section 3.1.

### 2.4 Feature Extraction

From the optimal correlation metric and temporal resolution identified in section 2.3, connectivity matrices were generated for each sliding window. From every window-derived matrix, we extracted a comprehensive set of 26 complex network measures, following the methodology established in [11]. These measures were computed for three distinct configurations derived from the best representation: (i) the full scalp connectivity matrix, (ii) the left hemisphere matrix, and (iii) the right hemisphere matrix. This design allowed us to quantify both global and hemispheric network organization and to assess their temporal evolution across the experimental sessions.

Prior to graph construction, all matrices were binarized using a threshold of 0.5 after standardizing [46] the matrix values, in accordance with our previous methodological framework [11]. Values above the threshold were assigned one, and those below or equal to 0, resulting in undirected, unweighted adjacency matrices.

The following complex network measures were then computed: assortativity coefficient [47, 48], average degree of k-nearest neighbors (kNN) [49], average shortest path length (APL) [50], betweenness centrality (BC) [51], closeness centrality (CC) [52], complexity, density [53], diameter [54], eccentricity [55], eigenvector centrality (EC) [56], efficiency [57], entropy of the degree distribution (ED) [58], hub score [59], k-core [60, 61], mean degree [62], second moment of the degree distribution (SMD) [63], and transitivity [64, 65].

This study used newly established measures, as thoroughly outlined in [11], to determine the number of communities in a complex network. First, community detection algorithms encompassed fast greedy (FC) [66], label propagation (LPC) [67], edge betweenness (EBC) [68], infomap (IC) [69], leading eigenvector (LC) [70], spinglass (SPC) [71], and multilevel community identification (MC) [72] were used to determine the communities within a network. Then, the largest community within each network is selected, followed by the computation of the average path length within that community, resulting in a singular metric. For clarity and coherence, we extended the abbreviations by appending the letter ‘A’ (indicating average path length) to denote the corresponding approach, resulting in AFC, AIC, ALC, ALPC, AEBC, ASPC, and AMC.

To ensure compatibility with statistical analyses and to avoid incorrect inference due to non-independence of sliding windows, complex network features extracted from all window-derived matrices within a segment were averaged, yielding a single representative value per subject, treatment, and time point. This strategy allowed us to assess how complex network features of brain connectivity vary over time and at different scales, while providing a richer, multi-resolution understanding of brain dynamics under the influence of ayahuasca. In line with our previous work [15], we employed a statistical tool known as linear mixed models (LMMs) to evaluate distributional differences in complex network features. We applied LMMs because they account for repeated measurements within subjects across timepoints and hemispheres, providing more reliable inference than traditional tests that assume independence [73–76]. This approach allowed us to rigorously assess both hemispheric asymmetries and temporal dynamics in network topology while mitigating the risk of inflated Type I errors, false positives that occur when a statistical test incorrectly detects a significant effect that is not truly present, that can result from ignoring subject-level dependencies [76].

All 26 features were statistically tested using LMMs across time points and hemispheres, and only those showing statistically significant effects after correction for multiple comparisons are presented and discussed in subsection 3.2, which includes the results for the full, left, and right hemisphere configurations.

### 2.5 Interregional node-strength analysis

Guided by prior work, summarized in Table 1, showing prominent frontal/temporal involvement and characteristic posterior—left and right temporal—central couplings, we quantified inter-regional node strength to examine connectivity patterns at the level of anatomically defined regions. The 23 scalp electrodes were grouped as follows: Frontal–Left (Fp1, F3, F7), Frontal–Right (Fp2, F4, F8), Frontal–Midline (Fz), Central–Left (C3, T7), Central–Right (C4, T8), Central–Midline (Cz), Parietal–Left (P3, P7), Parietal–Right (P4, P8), Parietal–Midline (Pz), Occipital–Left (O1), Occipital–Right (O2), Occipital–Midline (Oz), Temporal–Left (FT9), Temporal–Right (FT10), and Frontopolar–Midline (FPz).

For each subject and timepoint (T1, T2, T3), we computed the mean absolute connectivity strength between all channel pairs belonging to two distinct regions *r*_1_ and *r*_2_ (see Equation 1). As noted earlier, recordings from the placebo group were retained only at P3 and were used exclusively within the machine-learning analysis; therefore, the inter-regional node-strength analysis focuses solely on the ayahuasca group.

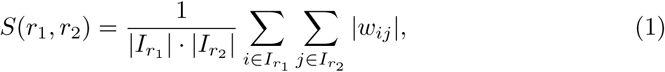

where *I_r__k_* denotes the set of channels assigned to region *r_k_*, and *w_ij_* represents the connectivity weight between channels *i* and *j*. In this context, inter-regional node strength reflects the mean of these connection weights, summarizing the overall intensity of functional coupling between regions. This procedure yielded a subject-level dataset of region-to-region strengths across time.

For statistical evaluation, we applied LMMs consistent with those described in the previous subsection 2.4, modeling timepoint (T1, T2, T3) as a fixed effect and subject as a random intercept. This design enabled us to assess longitudinal effects across all three timepoints, with specific pairwise contrasts estimated for T2 vs. T1, T3 vs. T1, and T3 vs. T2. Given the large number of region pairs, we controlled for multiple comparisons using the Benjamini–Hochberg procedure [77] with a false discovery rate (FDR) threshold of *q* = *α* = 0.05, ensuring that no more than 5% of the results declared significant are expected to be false positives [78–80].

The detailed findings of this inter-regional analysis, including statistically significant hemispheric and posterior–temporal–central interactions, are presented in subsection 3.3.

## 3 Results

### 3.1 ML results: Best representation selection

In figure S1 in supplementary material A, the performance of different correlation metrics (SC, PC, CCA, LW, TE) used to construct connectivity matrices is compared, reporting both training and independent test performance. SC achieved the highest and most stable results across all evaluation metrics (AUC, accuracy, recall, precision), followed closely by PC. LW performed moderately well, while CCA and TE showed weak discriminability, close to chance level. These results demonstrate that SC provided the most reliable representation of brain connectivity for distinguishing ayahuasca-induced brain states.

To evaluate the effect of temporal resolution, we compared classification performance across connectivity matrices derived from window sizes ranging from 10 to 120 s (Fig. S2 in supplementary material A). Performance increased steadily up to intermediate resolutions, with windows of 60–70 s yielding the highest and most stable results (test AUC ≈ 0.93, accuracy ≈ 0.93). Very short (10–20 s) and very long (110–120 s) windows produced weaker discriminability, suggesting that overly fine or overly coarse temporal resolutions obscure relevant connectivity patterns. Although the 60–70s provided the strongest classification, we retained the full range of window sizes for subsequent analyses to assess whether statistical results from complex network measures were consistent across temporal scales. This strategy ensured both methodological robustness and statistical validity by preventing biases related to the exclusive selection of a single window size.

For a state-comparison analysis, we fixed the representation to SC with 60-second windows—the best single configuration from the selection step—and compared pairs of conditions. As shown in Fig. S3 in supplementary material A, T1–T3 yielded the strongest separability (test *AUC* = 0.907, *accuracy* = 0.908, *recall* = 0.907, *precision* = 0.910), followed by T1 and T2 (test *AUC/accuracy* = 0.877). Discrimination between T2 and T3 was more modest (test *AUC* = 0.799). Importantly, a placebo vs. ayahuasca late comparison (P3–T3) still exceeded the 70% threshold (test *accuracy* = 0.750; *precision* = 0.833), indicating detectable group differences at the latest timepoint. This placebo comparison was used only for this sanity-check analysis; all remaining analyses exclude placebo and focus on ayahuasca time points.

To confirm that the observed separability was not specific to the SVM classifier, we fixed the representation to Spearman correlation (SC) with 60 s windows and focused on the T1–T2 comparison, which showed the most stable and well-balanced training and test performance. We then evaluated multiple machine learning models. As shown in Fig. S4 in supplementary material A, all classifiers—SVM, MLP, RF, and LR—achieved consistent test performance, with AUC and accuracy values exceeding 0.85. These convergent results demonstrate that the discriminative information captured by the selected representation is stable across different classification frameworks, supporting its robustness for downstream network analysis.

### 3.2 Complex network analysis results

We report complex-network features that demonstrated statistically robust effects, operationalized as LMM contrasts meeting the most stringent significance criterion (^∗∗∗^, *p <* 0.001) in at least one planned comparison. For each feature, an LMM was fitted with time point (T1, T2, T3) specified as a fixed effect and subject as a random intercept, from which the contrasts T2 vs. T1, T3 vs. T1, and T2 vs. T3 were derived.

As an initial step, we considered both hemispheres jointly by constructing connectivity networks from the complete set of electrodes. Complex-network features were extracted from these whole-brain representations and evaluated using the LMM framework. Notably, none of the features reached the most stringent significance threshold (^∗∗∗^, *p <* 0.001). A small number of features achieved moderate levels of significance (up to ^∗∗^, *p <* 0.01), primarily in the T2 vs. T1. These findings indicate that, when assessed at the level of both hemispheres combined, network-level alterations did not attain the robustness required for reporting under the strictest statistical criteria. For transparency, the complete set of moderate results is presented in Appendix B.

As illustrated in Fig. 2, eigenvector centrality (20 s window, Figure 2-a) revealed marked differences across time points. Both T2 and T3 showed significant decreases relative to T1, with p(T1 vs T2)=0.0060 and p(T1 vs T3)=0.0009. The average values decreased over time, with T2 and T3 showing lower centrality compared to T1, and T3 presenting the lowest values overall.

**Fig. 2.**
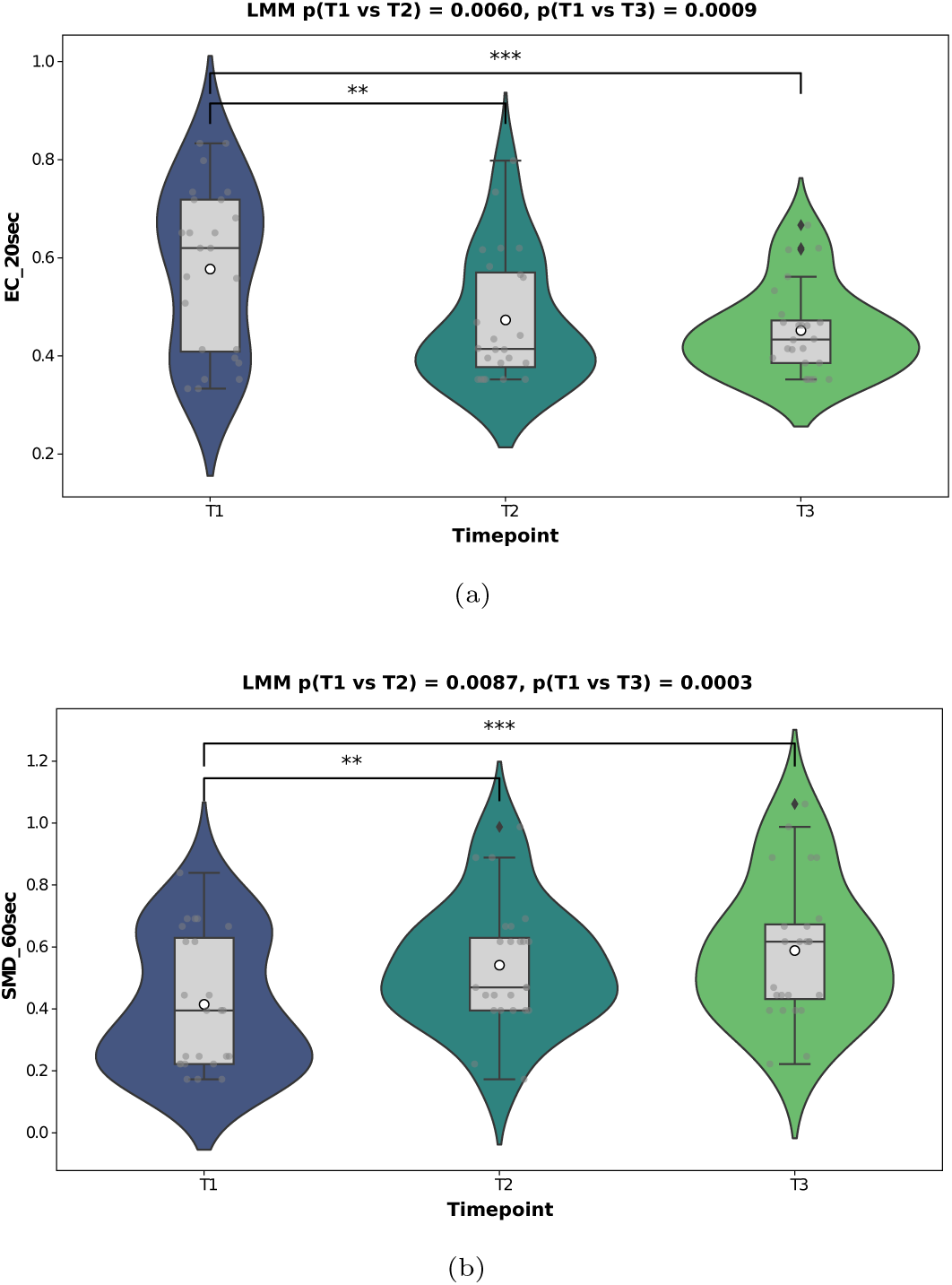
Rightelectrode complex network measures. (a) Eigenvector centrality (20s window): both T2 and T3 were significantly lower than T1, with *p*(*T* 1 vs *T* 2) = 0.0060 and *p*(*T* 1 vs *T* 3) = 0.0009; average values decreased across time, with T1 highest and T3 lowest. (b) Second moment of degree (60s window): both T2 and T3 were significantly higher than T1, with *p*(*T* 1 vs *T* 2) = 0.0087 and *p*(*T* 1 vs *T* 3) = 0.0003; average values increased across time, with T3 highest and T1 lowest. LMM *p*-values for the contrasts are shown in the panel titles. White dot = mean, box = interquartile range, whiskers = 1.5 IQR, outer shape = violin density. T1 = pre-dose (ayahuasca), T2 ≈ 2 h post-dose, T3 ≈ 4 h post-dose. **Significance:** ^∗∗∗^: *p <* 0.001, ^∗∗^: 0.001 *< p <* 0.01, ^∗^: 0.01 *< p <* 0.05, no star = *p* ≥ 0.05. **Abbreviations:** EC, eigenvector centrality; SMD, second moment of degree; LMM, linear mixed model.

Conversely, the SMD (60 s window, Figure 2-b) showed an opposite trend. Here, both T2 and T3 were significantly higher than T1, with p(T1 vs T2)=0.0087 and p(T1 vs T3)=0.0003. The metric increased across time, with T2 and T3 exhibiting higher values than T1, and T3 reaching the highest value overall. This divergence between the two metrics highlights complementary aspects of network topology: while eigenvector centrality reflects the global influence of highly connected hubs, the second moment quantifies degree heterogeneity, which is how unevenly connections are distributed across nodes.

Together, these complementary metrics reveal that ayahuasca induces both a reduction in hub dominance (eigenvector centrality) and an expansion of degree variability (SMD), offering convergent evidence of altered network organization across time.

As shown in Fig. 3, eigenvector centrality on the left hemisphere (Fig. 3-a and b) consistently revealed strong differences across time windows of 10 and 60 s. In both cases, T2 and T3 were significantly lower than T1, with p(T1 vs T2)=0.0009 and p(T1 vs T3)=0.0061 for the 10 s window, and p(T1 vs T2)=0.0005 and p(T1 vs T3)=0.0011 for the 60 s window. The average values decreased across time, with T2 and T3 lower than T1, and T3 presenting the lowest centrality overall.

**Fig. 3.**
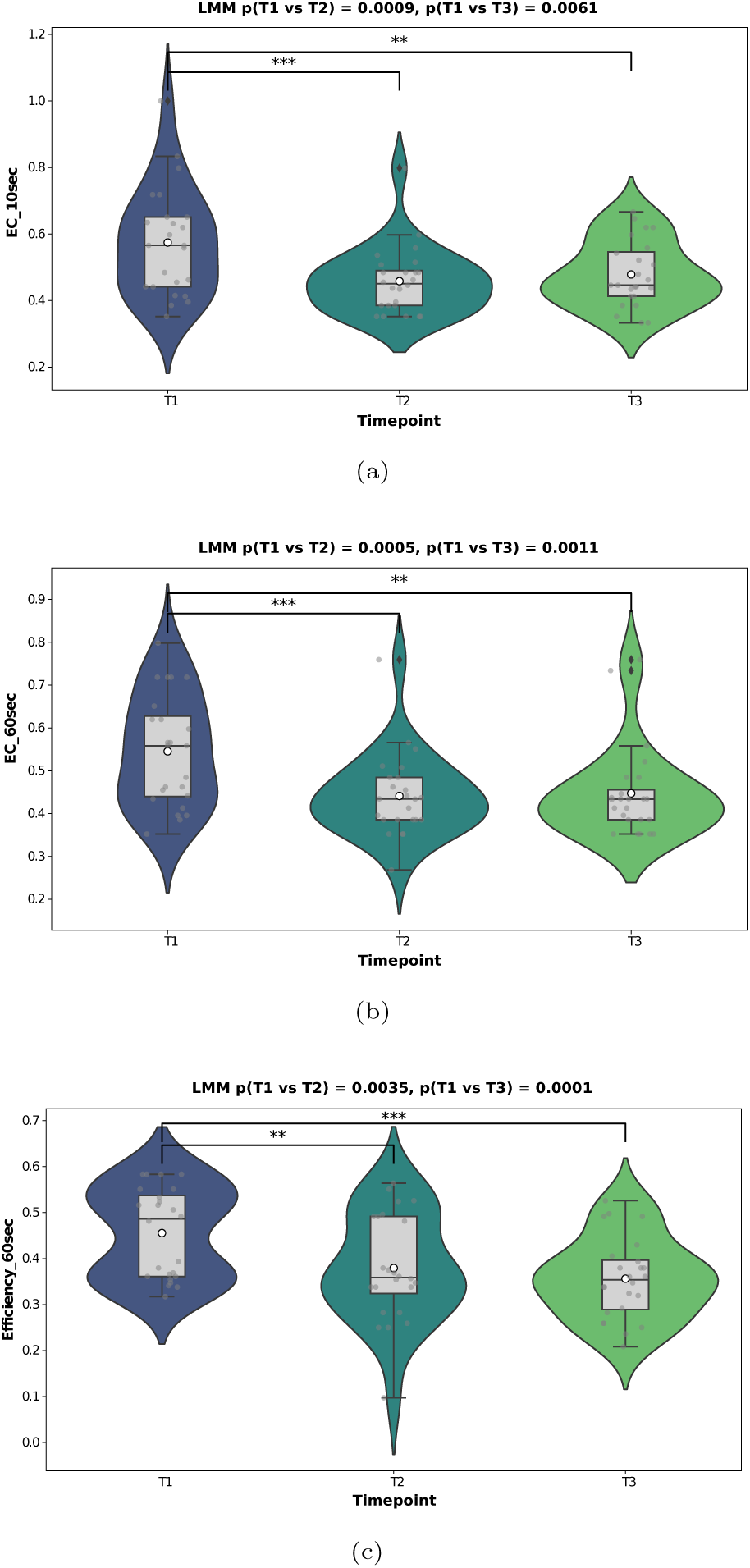
Left-electrode complex network measures. (a) Eigenvector centrality (10 s window) and (b) eigenvector centrality (60 s window): both T2 and T3 were significantly lower than T1, with *p*(*T* 1 vs *T* 2) = 0.0009 and *p*(*T* 1 vs *T* 3) = 0.0061 for the 10 s window, and *p*(*T* 1 vs *T* 2) = 0.0005 and *p*(*T* 1 vs *T* 3) = 0.0011 for the 60 s window; average values decreased with T1 highest and T3 lowest. (c) Global efficiency (60 s window): both T2 and T3 were significantly lower than T1, with *p*(*T* 1 vs *T* 2) = 0.0035 and *p*(*T* 1 vs *T* 3) = 0.0001; values were lower for T2 and T3 compared to T1, with T3 lowest. LMM *p*-values for the contrasts are shown in the panel titles. White dot = mean, box = interquartile range, whiskers = 1.5 IQR, outer shape = violin density. T1 = pre-dose (ayahuasca), T2 ≈ 2 h post-dose, T3 ≈ 4 h post-dose. **Significance:** ^∗∗∗^: *p <* 0.001, ^∗∗^: 0.001 *< p <* 0.01, ^∗^: 0.01 *< p <* 0.05, no star = *p* ≥ 0.05. **Abbreviations:** EC, eigenvector centrality; LMM, linear mixed model.

Global efficiency (Fig. 3-c, 60 s window) showed a similar decline at later time points. Here, both T2 and T3 were significantly lower than T1, with p(T1 vs T2)=0.0035 and p(T1 vs T3)=0.0001. Both T2 and T3 displayed lower efficiency compared to T1, with T3 showing the lowest values.

Overall, the observed alterations in network topology indicate both shared and lateralized components. Bilaterally, eigenvector centrality decreased from T1 to T3, suggesting a progressive weakening of hub influence over time. On the left hemisphere, this was accompanied by a reduction in global efficiency at 60s, consistent with less efficient large-scale integration. On the right hemisphere, by contrast, the most prominent change was an increase in degree dispersion (second moment of degree) rather than a further drop in efficiency, pointing to a more heterogeneous distribution of connectivity strengths. Notably, many of the strongest effects emerged at 60-second windows, which is the same temporal scale identified by the machine-learning framework as the most informative representation (Fig. S2). This convergence suggests that the 60-second window captures a biologically meaningful timescale of network reorganization under ayahuasca: long enough to integrate short-lived fluctuations in connectivity, yet short enough to resolve time-dependent changes in brain coordination.

To further assess hemispheric asymmetry, we performed direct statistical comparisons between left– and right-hemisphere electrode networks (Fig. 4). At the T3 timepoint, two metrics reached the significance level of 3 stars: the second moment of the degree (20 s window; Fig. 4-a) and network complexity (20 s window; Fig. 4-b). In both cases, the left hemisphere showed lower average values compared to the right, indicating that ayahuasca induces a lateralized effect at later stages.

**Fig. 4.**
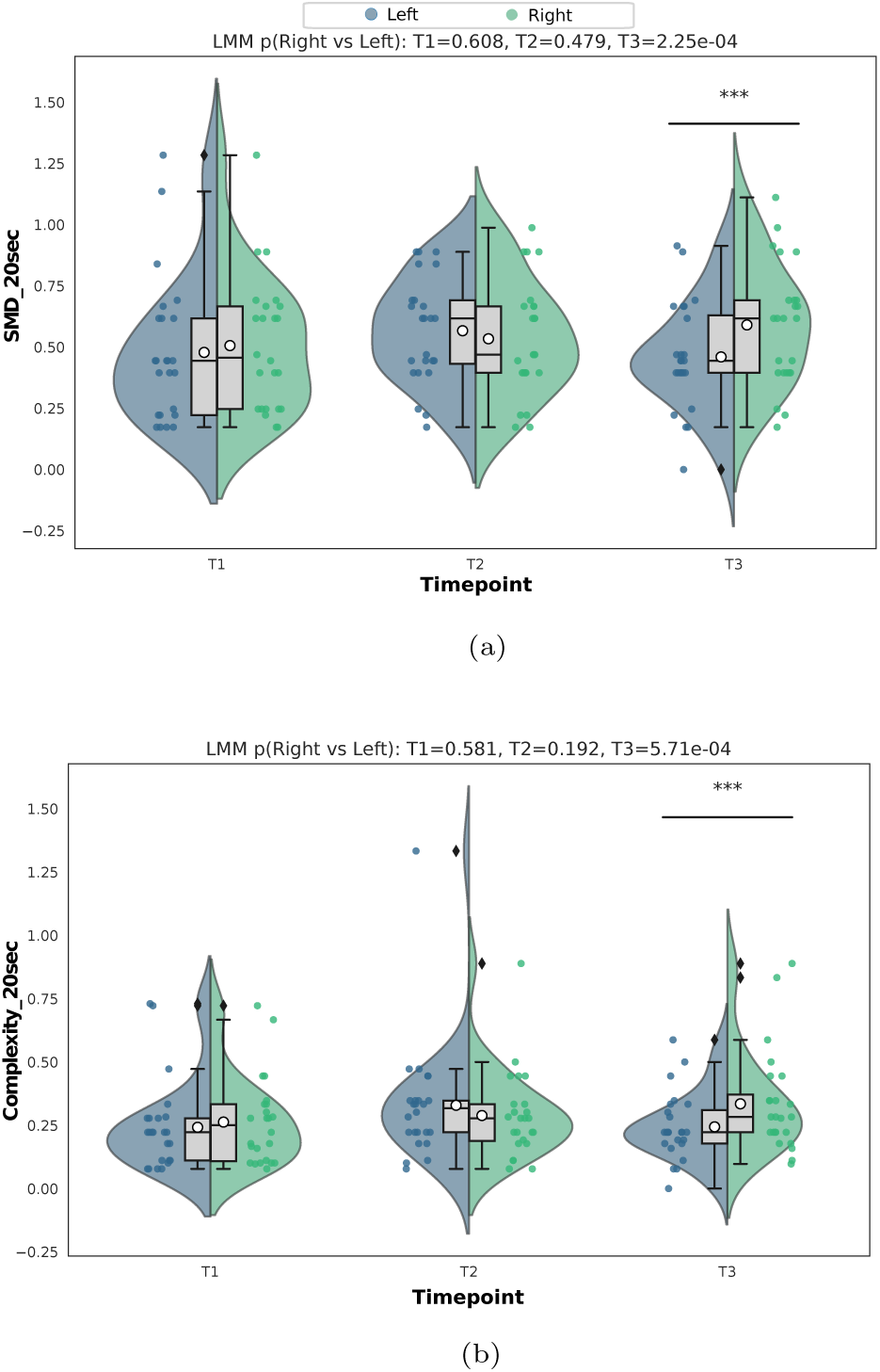
Hemispheric asymmetry in network topology. Direct comparisons between the left (blue) and right (green) hemispheres revealed significant differences at T3 in (a) the second moment of degree (20s window) and (b) complexity (20s window). In both metrics, the left hemisphere exhibited lower average values than the right. White dot = mean, box = interquartile range, whiskers = 1.5 IQR, outer shape = violin density. T1 = pre-dose (ayahuasca), T2 ≈ 2 h post-dose, T3 ≈ 4 h post-dose. **Significance:** ^∗∗∗^: *p <* 0.001, ^∗∗^: 0.001 *< p <* 0.01, ^∗^: 0.01 *< p <* 0.05, no star = *p* ≥ 0.05. **Abbreviations:** SMD, second moment of degree.

### 3.3 Interregional nodestrength results

The electrode pairs analyzed were derived from the anatomically defined regional grouping described in section 2.5, focusing on combinations that encompass the major posterior (occipital–temporal–parietal) and central pathways known to be modulated under psychedelics (Fig. 5). For the Occipital-Left-Parietal-Left connection, significant differences were observed between T1 and T2 (FDR-corrected *p* = 3.18 × 10^−5^) and between T1 and T3 (*p* = 2.80 × 10^−4^), with lower strength values at T2 and T3 and T2 showing the lowest (acute effect), while the contrast T2 vs. T3 was not significant (*p* = 0.920). A similar pattern was found for the Occipital-Left-Temporal-Left and Occipital-Left-Temporal-Right connections: both showed robust differences between T1 and T2 (FDR-corrected *p* = 1.76 × 10^−4^ and *p* = 2.09 × 10^−4^, respectively) and weaker, but still significant, differences between T1 and T3 (*p* = 0.018 and *p* = 0.019, respectively), whereas T2 vs. T3 remained non-significant in both cases (*p* = 0.808); again, T2 presented the lowest values. In contrast, the Temporal–Right–Central–Right connection displayed higher strength at T2 and T3, with a significant difference between T1 and T2 (FDR-corrected *p* = 1.76 × 10^−4^), but no significant difference between T1 and T3 (*p* = 0.081) or between T2 and T3 (*p* = 0.808); here, T2 showed the highest (acute) value. To facilitate interpretation, we additionally provide a schematic overview of the analyzed inter-electrode connections (Fig. 6). This visualization highlights that the Occipital-Left region participates in the largest number of connections (I-III), while the Temporal-Right region contributes to two of them (III and IV), underscoring their central roles in the analyzed network configuration.

**Fig. 5.**
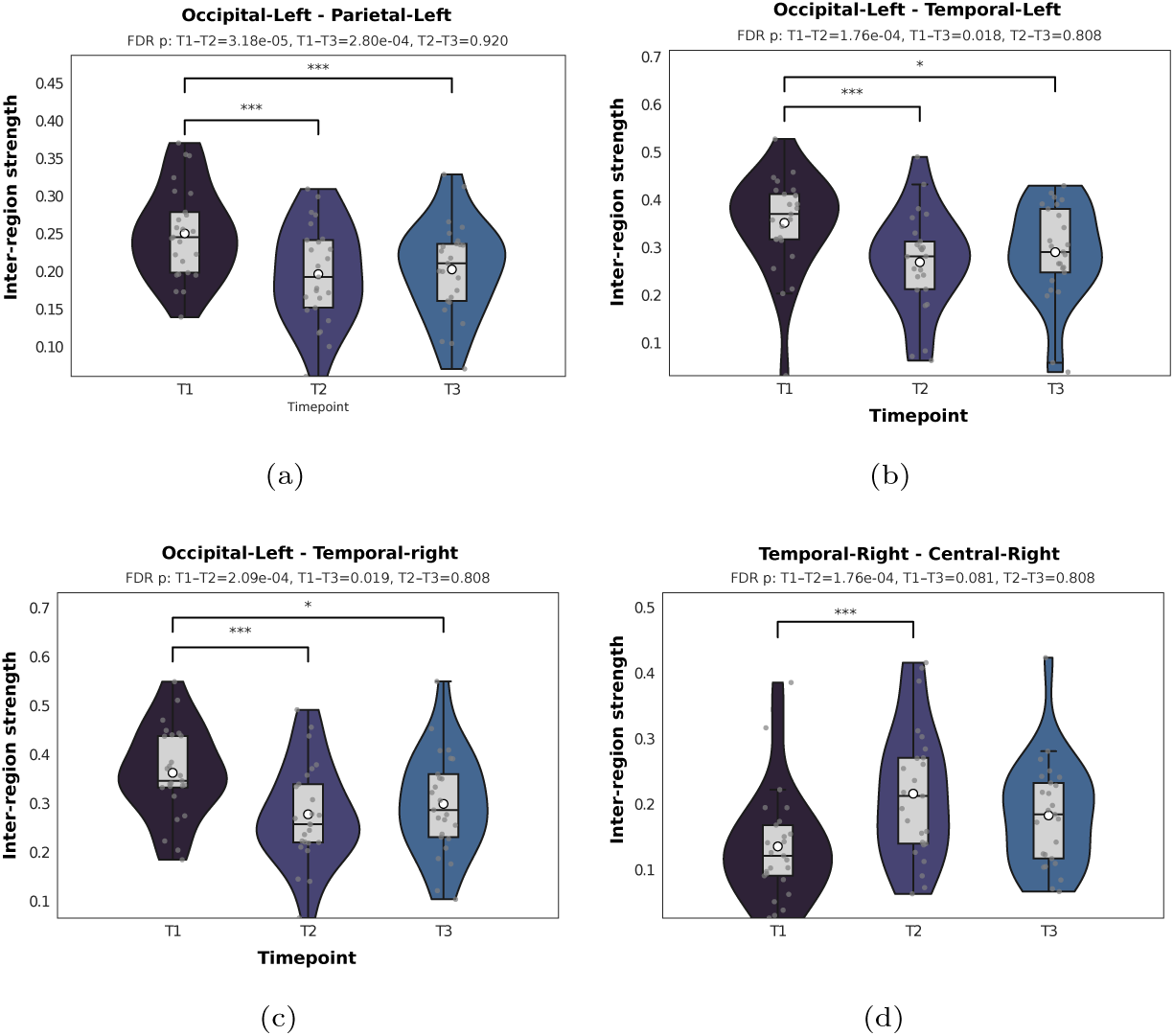
Interelectrode networks. Interregional strength comparisons for selected electrode pairs across T1–T3 are defined in Eq. 1. Panel titles display the region pair and FDR-corrected *p*-values from the Benjamini–Hochberg procedure. (a) Occipital–Left and Parietal–Left: significant differences for T1 vs. T2 (*p* = 3.18 × 10^−5^) and T1 vs. T3 (*p* = 2.80 × 10^−4^); T2 showed the lowest values. (b) Occipital–Left and Temporal–Left: T1 vs. T2 (*p* = 1.76 × 10^−4^) and T1 vs. T3 (*p* = 0.018) were significant; T2 showed the lowest values. (c) Occipital–Left and Temporal–Right: T1 vs. T2 (*p* = 2.09 × 10^−4^) and T1 vs. T3 (*p* = 0.019) were significant; T2 showed the lowest values. (d) Temporal–Right and Central–Right: T1 vs. T2 was significant (*p* = 1.76 × 10^−4^), whereas T1 vs. T3 was not (*p* = 0.081); T2 showed the highest values. White dot = mean, box = interquartile range, whiskers = 1.5 IQR, outer shape = violin density. T1 = pre-dose (ayahuasca), T2 ≈ 2 h post-dose, T3 ≈ 4 h post-dose. **Significance:** ^∗∗∗^: *p <* 0.001, ^∗∗^: 0.001 *< p <* 0.01, ^∗^: 0.01 *< p <* 0.05, no star = *p* ≥ 0.05. **Abbreviation:** FDR, false discovery rate.

**Fig. 6.**
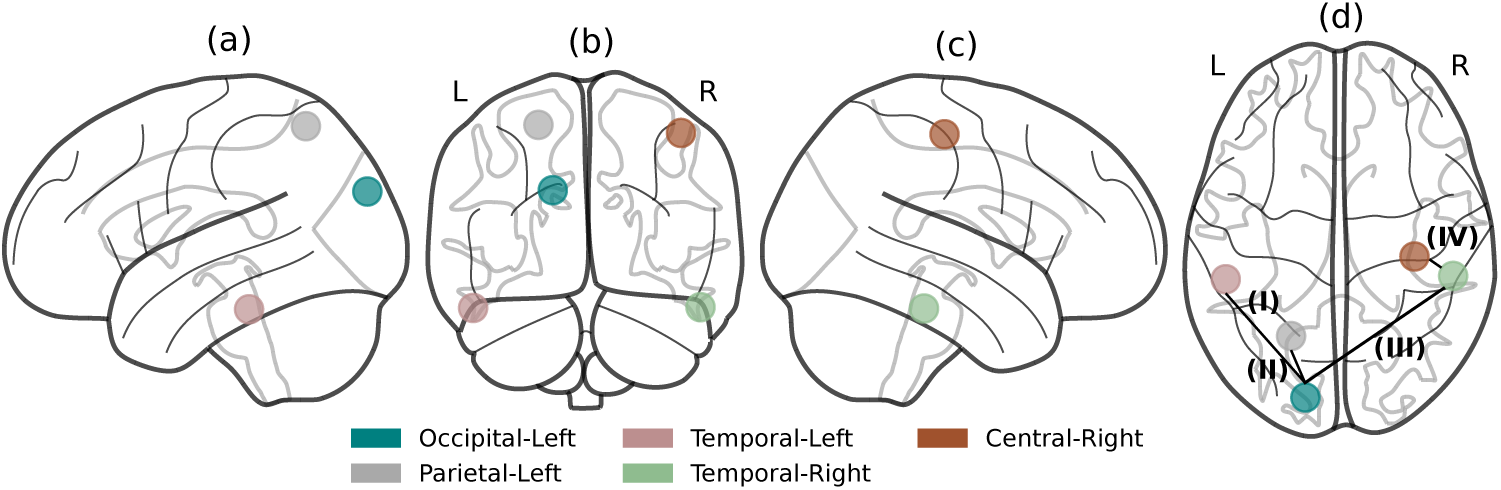
Summary of analyzed interelectrode connections. Brain schematic illustrating the five regions of interest and their pairwise connections from different anatomical perspectives (a–d). Colors indicate the regions: Occipital–Left (blue), Parietal–Left (gray), Temporal–Left (pink), Temporal–Right (green), and Central–Right (brown). Connections are numbered for clarity: (I) Occipital–Left with Temporal–Left, (II) Occipital–Left with Parietal–Left, (III) Occipital–Left with Temporal–Right, and (IV) Temporal–Right with Central–Right. These correspond to the statistical analyses shown in Fig. 5. The figure was elaborated by the authors using the Nilearn Python library [81].

## 4 Discussion

Among the correlation metrics tested to identify the most suitable approach for capturing brain changes, Spearman correlation yielded the most reliable connectivity representation (Fig. S1). Its non-parametric, rank-based nature is well suited to detect the increased complexity and non-Gaussian dependencies [82, 83] characteristic of psychedelic states, consistent with the entropic brain hypothesis described in Section 1 [3].

A central novelty of this study lies in the systematic evaluation of multiple temporal resolutions for connectivity estimation. From each 5-minute EEG segment, we generated connectivity matrices using sliding windows ranging from 10 to 120 seconds, in addition to the full segment, and averaged features across windows within each segment to ensure statistical validity. Machine learning analyses demonstrated that classification performance increased steadily with window size, peaking at inter-mediate resolutions of 60–70 seconds (test AUC = 0.93, accuracy = 0.93; Fig. S2). Notably, many of the significant effects in complex network measures also emerged at 60-second windows, mirroring the optimal resolution identified by the ML frame-work. This convergence highlights the importance of considering temporal resolution, as overly fine (10–20 s) or overly coarse (110–120 s) windows may obscure meaningful patterns of connectivity. Therefore, this indicates that temporal resolution is biologically meaningful, with the 60s scale capturing slower network reorganization under ayahuasca that is not as visible at shorter windows. By integrating results across multiple scales, our approach not only enhances methodological robustness but also reveals that intermediate temporal resolutions are biologically informative for capturing network reorganization under ayahuasca.

To further interrogate state-dependent effects, we fixed the representation to Spearman correlation with 60-second windows—the configuration identified as optimal in the selection step—and compared pairs of conditions (Fig. S3). Consistent with the statistical analyses, the highest discriminability was observed between T1 and T3 (test AUC/accuracy ≈ 0.91), followed by T1 and T2 (≈ 0.88). In contrast, T2–T3 yielded weaker performance (≈ 0.80), reflecting the absence of significant differences between these later states in the statistical tests. Together, these findings demonstrate that the most robust state contrasts occur between the early and later phases of ayahuasca, with intermediate-to-late transitions showing weaker separability, suggesting a progressive reorganization of connectivity rather than abrupt changes. Beyond identifying the optimal configuration to capture connectivity differences, the machine learning analysis also validated the statistical results of complex network measures, reinforcing that T1–T2 and T1–T3 differences are the most prominent, whereas T2–T3 transitions are less pronounced.

As an initial step, we evaluated global complex-network features derived from whole-brain connectivity networks constructed across all electrodes. Although none of these features reached the most stringent significance threshold (*p <* 0.001), and only a few achieved moderate significance (0.001 *< p <* 0.01), the direction of the effects suggested a more acute pattern of change at the whole-electrode level.

In the right-hemisphere networks (Figure 2), complex network analysis revealed marked temporal changes. Eigenvector centrality (EC, 20 s window, Figure 2-a) showed significant differences according to the LMM analysis, with p(T1 vs T2)=0.0060 and p(T1 vs T3)=0.0009. EC decreased progressively across time points, with values highest at baseline (T1, pre-dose), lower at T2 (2 h post-dose), and reaching their minimum at T3 (4 h post-dose), indicating a decline in hub prominence and a redistribution of influence across the network over time.

Conversely, the second moment of the degree distribution (60 s window, Figure 2-b) showed the opposite temporal pattern. This metric increased over time, with T2 and T3 being higher than T1, and T3 having the highest value overall. LMM contrasts indicated significant differences for both p(T1 vs T2)=0.0087 and p(T1 vs T3)=0.0003. Mechanistically, a rise in degree heterogeneity is compatible with the recruitment of alternative pathways: as hub influence declines, previously weak or cross-module connections may strengthen or become supra-threshold, effectively introducing additional routes for information flow.

Consistently, in the left hemisphere (Figure 3), EC at both 10 s and 60 s windows (Figure 3-a,b) showed strong temporal differences with the same monotonic decrease (T1 highest; T2 and T3 lower; T3 lowest). For EC at 10 s, the contrasts yielded p(T1 vs T2) = 0.0009 and p(T1 vs T3) = 0.0061. For EC at 60 s, the results show p(T1 vs T2) = 0.0005 and p(T1 vs T3) = 0.0011. On the left hemisphere, global efficiency (60 s window; Figure 3-c) also declined at later time points, with significant differences of p(T1 vs T2)=0.0035 and p(T1 vs T3)=0.0001. Global efficiency indexes the ease of network-wide communication—higher values reflect shorter average topological paths and greater integration—so the efficiency drop suggests that as hub-mediated shortcuts contribute less, average paths lengthen and communication increasingly relies on alternative but less efficient routes. Notably, many of the strongest effects emerged at 60 s, the same temporal scale identified by the machine-learning framework as most informative (Figure S2), reinforcing that this timescale captures slower network reorganization under ayahuasca. As outlined in Section 1 and Table 1, convergent neuroimaging across serotonergic psychedelics points to attenuated hub dominance and reduced within-network integrity—especially within the DMN. Furthermore, to support our results, the same reduction in within-network integrity can be found in other psychedelics like lysergic acid diethylamide (LSD)[84–87]. For ayahuasca specifically, reduced PCC/pre-cuneus connectivity has been reported in [2], and our prior EEG study observed slower network-wide accessibility (lower closeness centrality) [11]. In this context, the bilateral EC decreases and the left-hemisphere drop in global efficiency observed here index a loss of hub-centric control and diminished largescale integration at the sensor level, while the increase in degree heterogeneity (*right > left*), operationalized as a rise in the second moment of the degree distribution and corroborated by higher complexity, is compatible with recruitment of alternative, less efficient routes. Taken together, these features align with increased cross-network coupling under psychedelics [84–86, 88, 89] and with the less constrained, more entropic regime introduced in Section 1.

To assess hemispheric asymmetry, we conducted direct statistical comparisons between left– and right-hemisphere electrode networks (Figure 4). At T3, two metrics showed robust hemispheric differences: the second moment of the degree (20,s window; LMM *p* = 2.25 × 10^−4^) and network complexity (20,s window; LMM *p* = 5.71 × 10^−4^). In both cases, the left hemisphere exhibited lower average values than the right, indicating a lateralized effect emerging at later stages under ayahuasca. At T1 and T2, no significant hemispheric differences were detected for these metrics (all *p >* 0.05). Here, we define complexity as the variance of the node-degree distribution normalized by the mean degree (see Eq. 2):

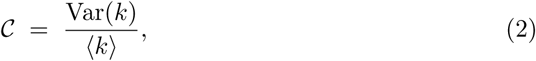

where *k* = (*k*_1_*, …, k_N_*) are the node degrees (loops permitted), 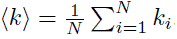, and Var(*k*) is the variance of the node degrees.

Accordingly, at T3, both the second moment of degree and the complexity metric were higher in the right hemisphere than in the left, indicating a more heterogeneous and dispersed degree structure on the right. In other words, node degree values are more unevenly distributed in right-hemisphere networks at the late time point. Combined with the bilateral decrease in eigenvector centrality and the left-hemisphere drop in global efficiency at 60s, this pattern suggests that hub-mediated shortcuts weaken overall, while the right hemisphere develops a more heterogeneous connectivity profile. Rather than implying a uniform increase in the number of right-hemisphere connections, these findings are more consistent with a right-lateralized shift toward more variable and distributed connectivity patterns.

When analyzing both hemispheres together, effects appeared less pronounced than in hemisphere-specific analyses. One interpretation is that hemispheric compensation or balancing mechanisms dampen unilateral alterations when networks are aggregated. Extensive evidence shows that cross-hemispheric coordination can buffer unilateral reorganization where contralateral circuits dynamically compensate for local disruptions, thereby stabilizing global functional topology and masking hemisphere-specific effects at the whole-brain level [90–93].

Finally, pairwise node-strength analyses (Figure 5) reveal a clear posterior–left attenuation: Occipital-Left × Parietal-Left, Occipital-Left × Temporal-Left, and Occipital-Left × Temporal-Right all decrease, with the sharpest drop at the inter-mediate post-dose state. In sensor-space terms, these links implicate visual (O1) and posterior association territories (P3/P7; FT9/FT10) and are topographically consistent with reduced posterior midline influence (PCC/precuneus) within the DMN noted in prior ayahuasca work. This regional pattern aligns with the bilat-eral eigenvector-centrality decrease and the left-hemisphere efficiency drop (reduced large-scale integration) reported here, cohering with the DMN findings summarized in Table 1 and described in the Section 1.

By contrast, a right-lateralized reinforcement emerges for Temporal-Right × Central-Right (FT10 × C4/T8), suggesting a transient reweighting of right-hemisphere pathways that is compatible with salience– and attention-related circuitry described in Section 1. Coupled with a right *>* left rise in degree dispersion (second moment and complexity), these effects suggest that, as posterior left-hemisphere connections weaken, some communication is supported by more heterogeneous and distributed routes that are more prominently expressed in the right hemisphere. This pattern reflects a partial rerouting of information flow rather than a global increase in right-hemisphere connectivity and mirrors the broader DMN/SN modulation and entropy-related shifts reviewed in Table 1.

The hemispheric asymmetry observed here also resonates with conditions characterized by atypical lateralization patterns. In autism spectrum disorder, several studies, including our recent work [94–96], have reported stronger left-hemisphere connectivity or reduced right-hemisphere integration compared with neurotypical controls. Although the present findings were obtained in healthy, naïve ayahuasca users and cannot be extrapolated to clinical settings, they show that ayahuasca can selectively shift hemispheric balance in large-scale EEG-derived networks. This highlights a potential direction for future mechanistic research: examining whether ayahuasca-induced reorganization of lateralized connectivity may help elucidate how atypical hemispheric patterns arise or are compensated for in neurodevelopmental conditions. Such considerations remain strictly conceptual and non-therapeutic, but they indicate a promising avenue for comparative studies across populations with distinct lateralization profiles.

### 4.1 Novelty and methodological advance

Novelty and methodological advance. Building on our prior EEG study [11], we extend the pipeline by (i) temporal-resolution triangulation (systematically sweeping 10–120 s windows) and showing convergence between LMM effects and ML separability at 60 s; (ii) hemisphere-resolved, dispersion-sensitive topology (eigenvector centrality, second moment of degree, and an interpretable complexity index Var(k)/〈k〉); and (iii) edge-level mapping via inter-regional node strength. All complex-network measures were computed within sliding windows and then averaged per 5-min segment to stabilize estimates and ensure fair comparisons across hemispheres and time points. This integrated approach reveals a right-lateralized increase in degree dispersion at T3 and a left-hemisphere efficiency drop at 60 s, with the same 60 s window emerging as ML-optimal—evidence that temporal scale is biologically meaningful in capturing ayahuasca-induced network reorganization.

Third, we extend the analysis to edge-level mapping via interregional node strength, demonstrating a clear posterior–left attenuation together with a transient strengthening of right temporal–central coupling. Notably, this pattern parallels our previous findings in the DMT cohort [6], where we also reported concurrent decreases in integration and segregation alongside the emergence of alternative communication routes.

Together, these convergent findings update the mechanistic interpretation of psychedelic network dynamics. The bilateral loss of integration echoes the reduced segregation/integration balance observed under LSD [88] and supports the entropic-brain framework [97]. At the same time, the localized reinforcement of right temporal–central pathways aligns with the view articulated in [98] that psychedelics do not merely increase randomness: disruption of normative architecture can coexist with the appearance of strong, topologically far-reaching connections. By integrating temporal-scale optimization, hemispheric topology, and edge-level mapping—and by extending pat-terns previously observed under DMT [6]—our framework captures both aspects of psychedelic network reorganization: the weakening of canonical integration and the rerouting of communication through alternative pathways.

## 5 Conclusion

This study applied an integrated framework combining EEG connectivity, machine learning, and complex network analysis to investigate how ayahuasca modulates largescale brain organization over time in a cohort of naïve users, where neural effects are typically attenuated. By linking data-driven model selection with graph-theoretical interpretation, we provide a multilevel view of both temporal and hemispheric network reorganization. Using a pipeline that couples machine learning with complex-network analysis, we identified Spearman correlation as the most reliable connectivity representation and showed that a temporal scale of approximately 60 s is biologically informative, as it maximized classification performance and coincided with the strongest network effects. Across both hemispheres, eigenvector centrality decreased from T1 to T2 to T3, indicating weakening hub influence. In the right hemisphere, the second moment of degree increased at 60 s, while in the left hemisphere, global efficiency decreased at 60 s. Direct comparisons at T3 showed a higher degree of dispersion on the right (greater second moment and complexity), and edge-level analyses revealed acute posterior-left weakening with a transient strengthening between the right temporal and central regions. Taken together, these convergent findings support a concise interpretation: as hub-centric shortcuts weaken, communication is increasingly routed through alternative, more distributed—and less efficient—pathways, with a more heterogeneous and selectively strengthened pattern emerging in right-hemisphere networks at the later time point. The results extend our previous methodology by demonstrating how window-optimized representations, hemisphere-resolved dispersion metrics, and edge-wise validation can jointly expose psychedelic-induced reorganization of brain networks.

## Data Availability

Data analysed during this study are available from DBA upon reasonable request.

## Acknowledgements

We would like to thank Dr. Manuel Ciba for his valuable statistical suggestions, particularly regarding the implementation of the linear mixed model framework.

## A Supplementary Results: ML results

**Fig. S1.**
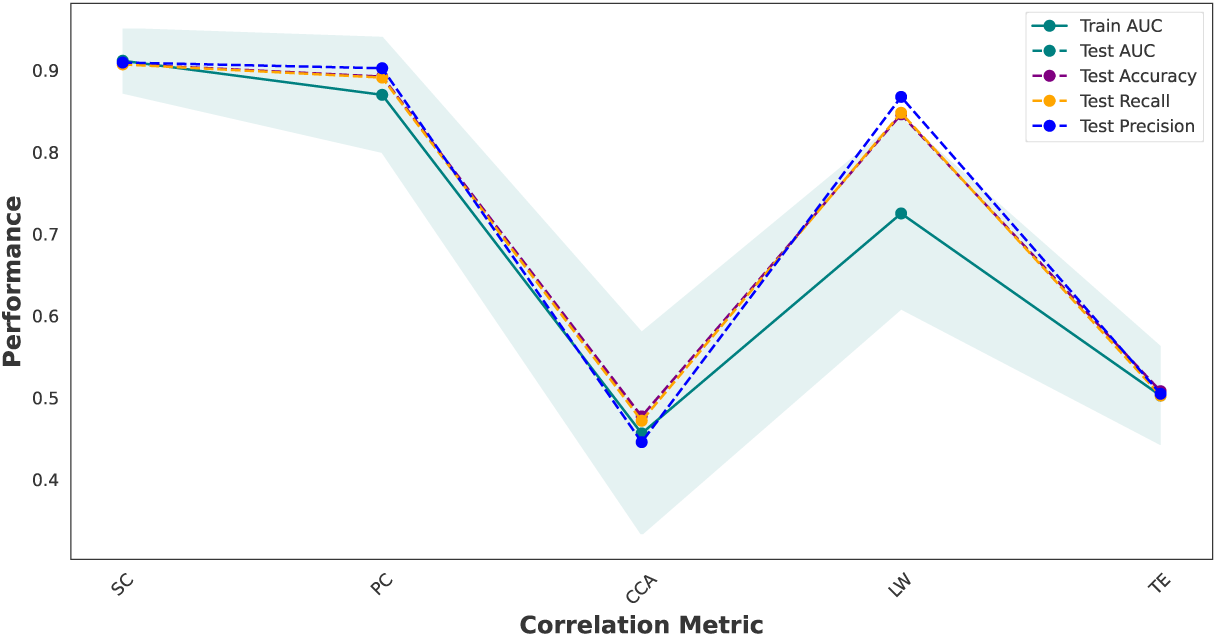
Performance across correlation metrics. Comparison of connectivity representations constructed using five correlation methods: SC, PC, CCA, LW, and TE. Classification was performed using an SVM. For the AUC metric, the solid green line represents the mean training AUC across folds, and the green shaded region shows the standard deviation of the training AUC. The dashed green line corresponds to the test AUC, but because training and test AUC values were nearly identical across all folds, the dashed green test curve is almost entirely overlapped by the solid green training curve and therefore not visually distinguishable in the figure. Furthermore, dashed lines indicate test performance accuracy (purple), recall (yellow), and precision (blue). SC achieved the highest overall performance. The dashed green line corresponds to the test AUC; however, train and test AUC values are almost identical, and therefore the dashed test curve is largely overlapped by the training curve and not easily visible in the figure. **Abbreviations:** AUC, area under the ROC curve; PC, Pearson correlation; SC, Spearman correlation; CCA, canonical correlation analysis; LW, Ledoit–Wolf shrinkage covariance; TE, transfer entropy; SVM, Support Vector Machine.

**Fig. S2.**
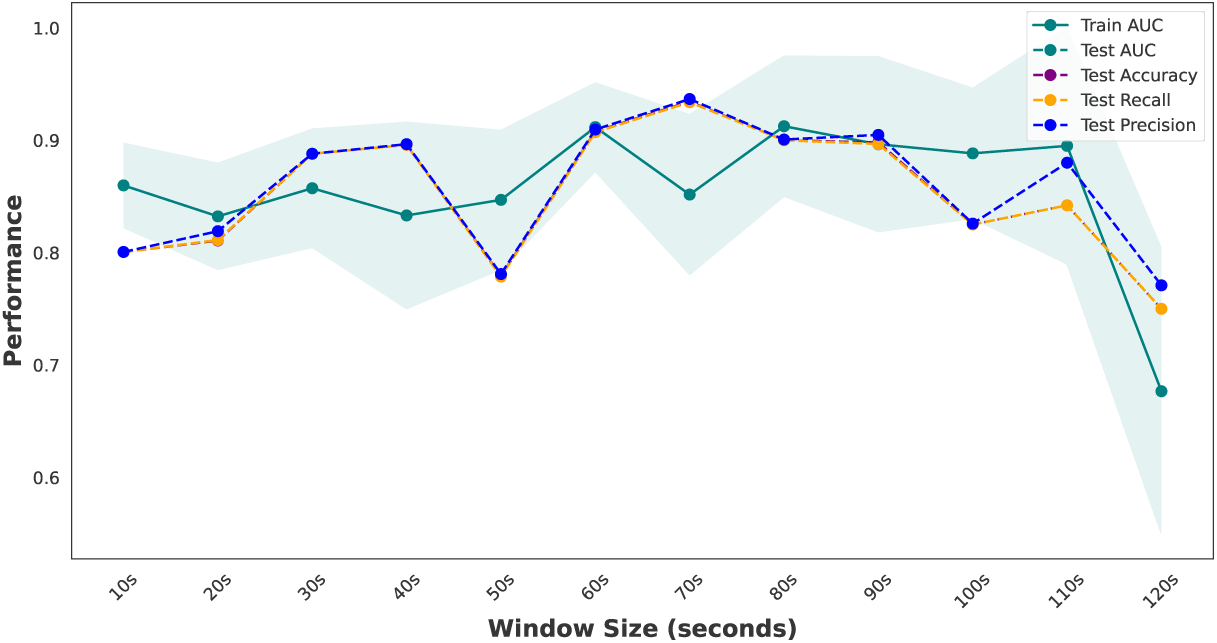
Performance across sliding-window sizes. Average classification metrics are shown for connectivity matrices computed with temporal windows from 10 to 120 s. For the AUC metric, the solid green line represents the mean training AUC across folds, and the green shaded region shows the standard deviation of the training AUC. The dashed green line corresponds to the test AUC, but because training and test AUC values were nearly identical across all folds, the dashed green test curve is almost entirely overlapped by the solid green training curve and therefore not visually distinguishable in the figure. Furthermore, dashed lines indicate test performance for accuracy (purple), recall (yellow), and precision (blue). The dashed green line corresponds to the test AUC; however, train and test AUC values are almost identical and therefore the dashed test curve is largely overlapped by the training curve and not easily visible in the figure. Windows of the 60–70s achieved the highest and most stable performance. **Abbreviations:** AUC, area under the ROC curve.

**Fig. S3.**
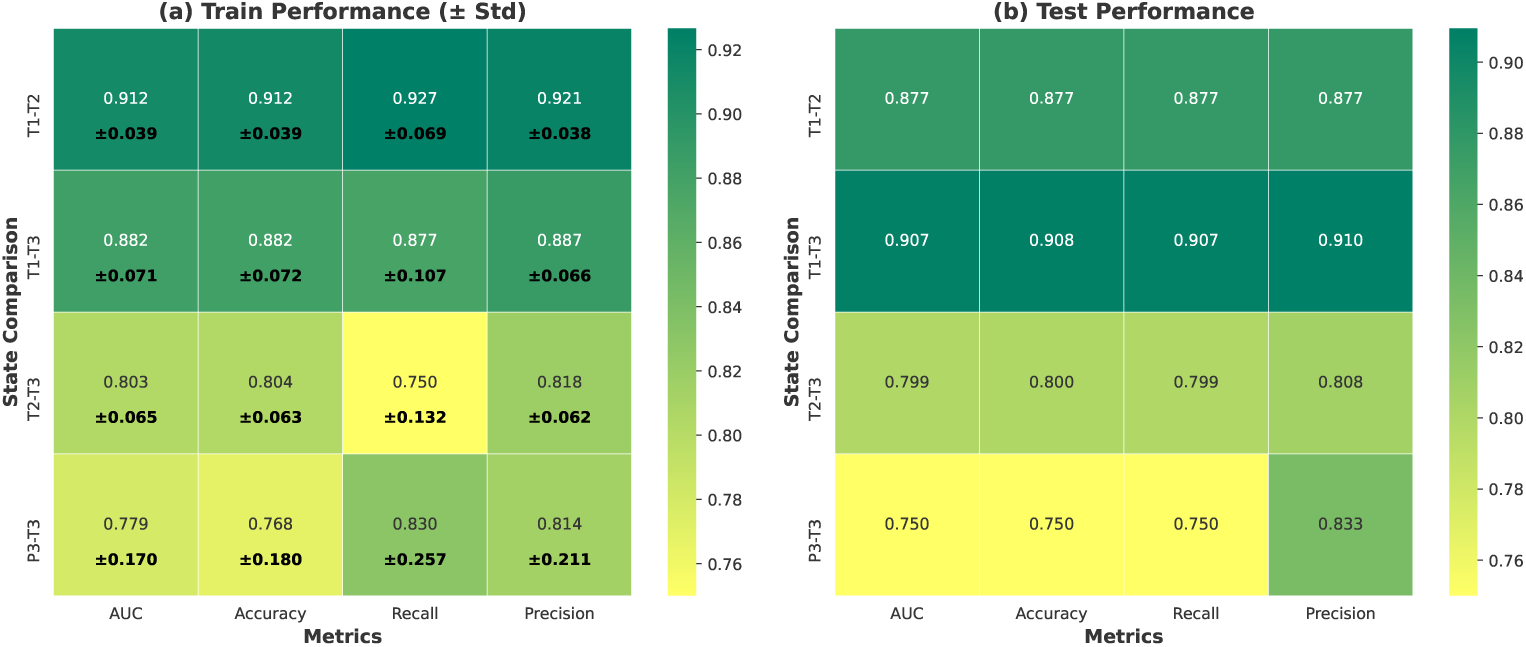
State-wise classification using the selected representation (SC, 60 s) with SVM. Heatmaps summarize (a) training performance (cells show means with the “±” standard deviations over folds overlaid in each cell) and (b) test performance for four pairwise comparisons: T1–T2, T1–T3, T2–T3, and P3–T3. T1 = pre-dose (ayahuasca), T2 ≈ 2 h post-dose, T3 ≈ 4 h post-dose; P3 = placebo at ≈ 4 h. T1–T3 showed the best separability (test AUC and accuracy ≈ 0.91), followed by T1–T2 (≈ 0.88), and T2–T3 was lower (≈ 0.80). T1–T3 showed slightly higher test performance but a mild drop in training accuracy, suggesting less consistent learning stability. The placebo vs. ayahuasca late comparison (P3–T3) exceeded the 70% threshold (test accuracy = 0.75, precision = 0.83), supporting detectable group differences at the late timepoint, indicating detectable group differences at the latest time point. **Abbreviations:** AUC, area under the ROC curve; SVM, Support Vector Machine.

**Fig. S4.**
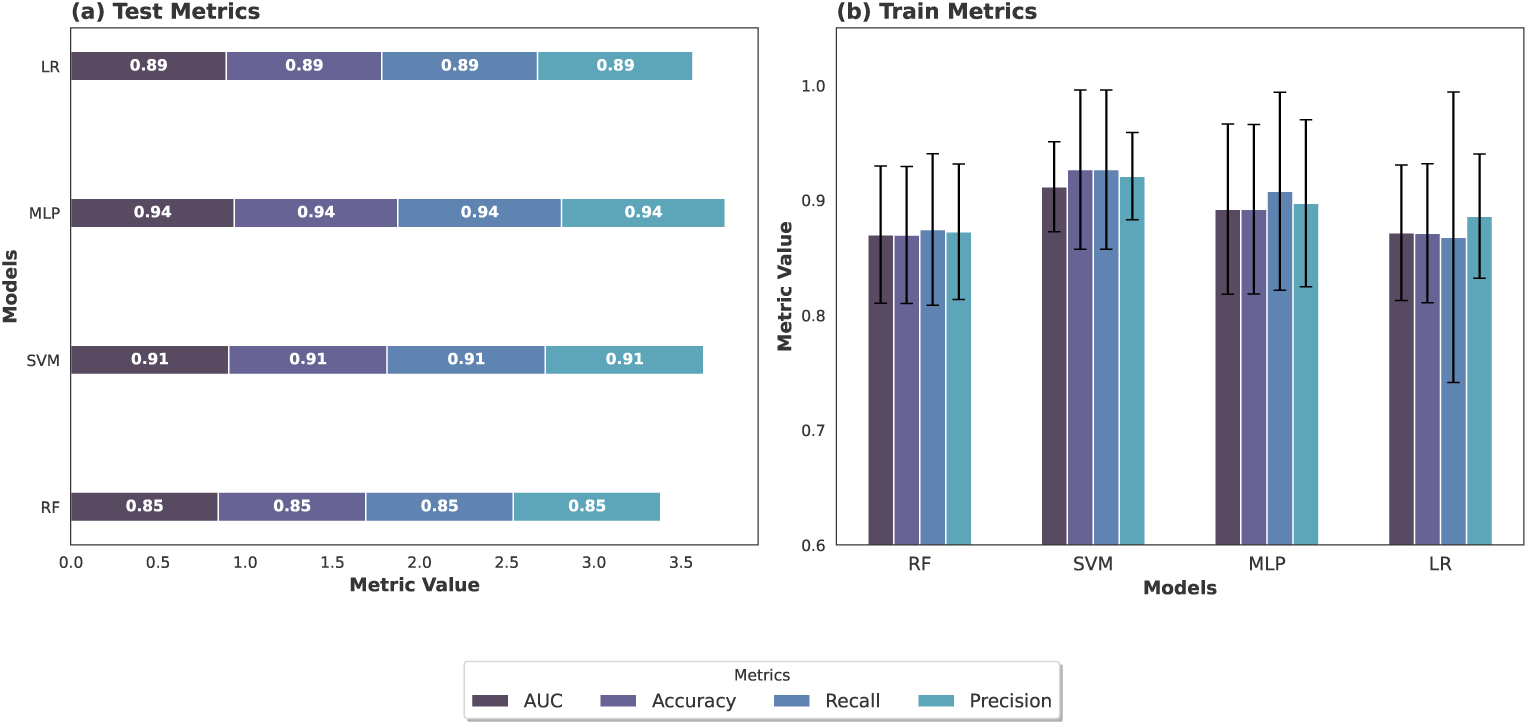
Validation across classifiers for the T1–T2 comparison. Using the connectivity representation (SC, 60 s windows), classification performance was evaluated across multiple algorithms: SVM, MLP, RF, and LR. Bar plots show mean test (a) and train (b) performance across AUC, accuracy, recall, and precision. All classifiers achieved comparably high results (AUC and accuracy *>* 0.85), confirming that the discriminability of the T1-T2 state comparison was not model-specific. T1 = pre-dose (ayahuasca), T2 ≈ 2 h post-dose. **Abbreviations:** AUC = area under the ROC curve; SVM = support vector machine; MLP = multilayer perceptron; RF = random forest; LR = logistic regression.

## B Supplementary Results: Moderate Effects from Whole-Electrode Networks

When considering the entire electrode set (both hemispheres jointly), no complex-network feature reached the most stringent significance threshold (^∗∗∗^, *p <* 0.001). However, several features exhibited moderate effects, attaining significance at the ^∗∗^ level (*p <* 0.01). For completeness, these results are presented here as supplementary material. Figure S5 illustrates the distribution of these features across timepoints (T1, T2, T3) using boxplots, with significance annotations corresponding to the LMM contrasts.

**Fig. S5.**
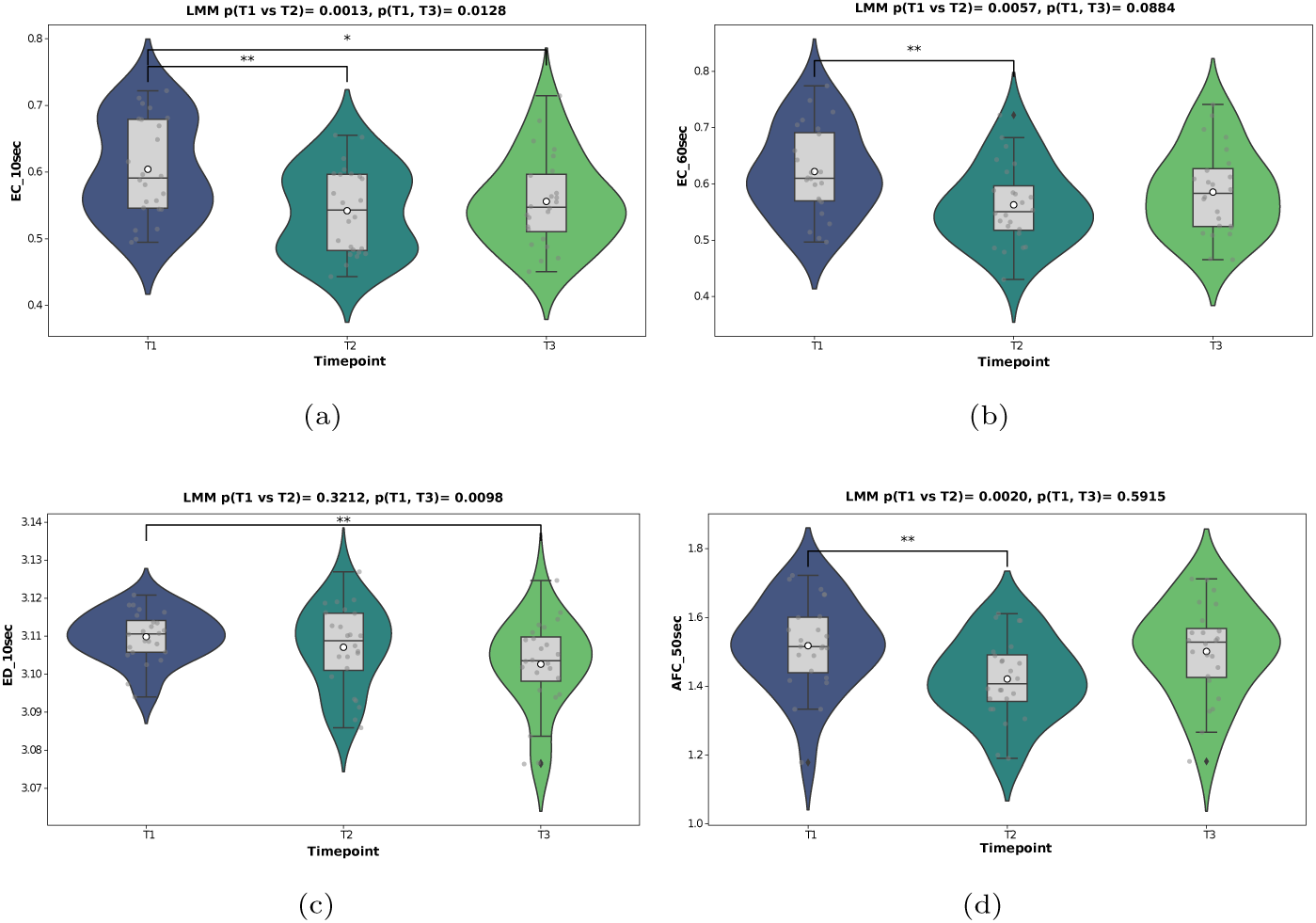
Moderate effects in whole-electrode networks. Examples of complex-network features that reached moderate significance (^∗∗^, *p <* 0.01) when derived from the full electrode set. Each panel shows the distribution of values across timepoints (T1, T2, T3) using boxplots, with significance annotations corresponding to the LMM contrasts. T1 = pre-dose (ayahuasca), T2 ≈ 2 h post-dose, T3 ≈ 4 h post-dose. LMM *p*-values for the reported contrasts are shown in the panel titles. **Abbreviations:** EC, eigenvector centrality; ED, entropy degree; AFC, average path length within the largest community detected by the Fast Greedy algorithm; LMM, linear mixed model.

